# Approximating conformational Boltzmann distributions with AlphaFold2 predictions

**DOI:** 10.1101/2023.08.06.552168

**Authors:** Benjamin P. Brown, Richard A. Stein, Jens Meiler, Hassane Mchaourab

## Abstract

Protein dynamics are intimately tied to biological function and can enable processes such as signal transduction, enzyme catalysis, and molecular recognition. The relative free energies of conformations that contribute to these functional equilibria are evolved for the physiology of the organism. Despite the importance of these equilibria for understanding biological function and developing treatments for disease, the computational and experimental methods capable of quantifying them are limited to systems of modest size. Here, we demonstrate that AlphaFold2 contact distance distributions can approximate conformational Boltzmann distributions, which we evaluate through examination of the joint probability distributions of inter-residue contact distances along functionally relevant collective variables of several protein systems. Further, we show that contact distance probability distributions generated by AlphaFold2 are sensitive to points mutations thus AF2 can predict the structural effects of mutations in some systems. We anticipate that our approach will be a valuable tool to model the thermodynamics of conformational changes in large biomolecular systems.

## Introduction

The presiding paradigm in protein structural biology is that amino acid sequence determines three-dimensional structure and that structure imparts function ^1,2^. Frequently, however, the functional properties of a protein are not solely determined by a single stable tertiary structure, but by the dynamics of conformational changes in that structure. This dynamic behavior can occur on a range of timescales, from nanoseconds to milliseconds or longer, and can involve local or global movements of a protein. Correspondingly, protein dynamics are involved in a range of biological processes, including folding, signal transduction, enzyme catalysis, and molecular recognition.

Methods of experimental structural biology, such as X-ray crystallography and cryogenic electron microscopy (cryo-EM), can visualize snapshots of these motions as changes in packing of secondary structure elements and/or the arrangement of individual folded domains. In actuality, these snapshots represent a subset of accessible states in the larger conformational free energy landscape of the system. The relative free energies of and rates of interconversion between these conformations control biological function. Consequently, perturbations to the chemical structure of a protein through missense mutations or truncations can cause disease by altering the relative energetics and kinetics of these conformational equilibria. Parsing the detailed mechanisms of protein conformational changes requires complementary use of structure determination tools with methods that provide insight into protein dynamics and/or state distributions.

For example, cancers caused by mutations to epidermal growth factor receptor (EGFR) are arguably some of the prominent examples of the relationship between aberrant protein dynamics and disease ^3–11^. Nuclear magnetic resonance (NMR) and other forms of spectroscopy have been applied to great effect to identify dynamic mechanisms of molecular recognition and function in proteins, e.g., kinases ^12–15^. We ^3–5^ and others ^6–11^ have employed molecular dynamics (MD) simulation approaches to probe the energetics of the EGFR kinase domain and its mutants. Despite the utility of such methods, solution NMR is limited to protein systems of modest sizes, and the computational cost of MD simulations at biologically relevant timescales of larger protein systems is prohibitive to all but the most specialized hardware, such as the Anton 2 supercomputer ^16^. Consequently, new scalable methods that can sample protein conformations and return a Boltzmann distribution of states are highly desirable.

Recently, the application of AlphaFold2 (AF2) and other protein structure prediction artificial intelligence (AI) models to the study of protein dynamics has transformed the field of computational structural biology. Perhaps most intuitively, conformers generated with AF2 have successfully been used to seed MD simulations and construct conformational free energy surfaces through Markov state modeling ^17^. Several strategies have been proposed that utilize machine learning (ML) to identify collective variables (CV) that can be used to bias simulations toward more frequent sampling of rare events ^18–22^. Indeed, learned contact distances have been directly employed as biasing potentials in enhanced sampling MD simulations ^23,24^. Still other MD simulation approaches apply ML to improve reconstruction of all-atom structures from coarse-grained (CG) representations or increase the accuracy of CG free energy landscapes^25,26^.

Distinct from these approaches are investigations into the extent to which protein structure prediction AI has directly learned to approximate conformational flexibility. For example, a few early studies have shown that AF2’s per-residue confidence score pLDDT correlates with flexibility metrics obtained from all-atom and coarse-grained MD simulations ^27,28^ and backbone NMR N-H S^2^ order parameters ^29^. Notably, these studies have restricted their focus to correlations between metrics of predicted structure fidelity (i.e., pLDDT and PAE) and time-averaged aggregated atomic fluctuations. It remains unclear whether protein structure prediction AI have learned thermodynamic information, especially as it relates to conformational free energies.

The question, “Can AI learn the thermodynamics of biomolecular conformational ensembles?” has potentially dramatic implications for the future of molecular modeling and simulation. Here, we investigate the extent to which the contact distance distributions generated by AF2 approximate conformational Boltzmann distributions for functionally relevant CVs and compare these distributions to those obtained by electron paramagnetic resonance (EPR) spectroscopy and MD simulations. We then apply AF2 to predict the thermodynamic consequences of mutations in several receptor tyrosine kinases. Overall, we conclude that AF2 can provide coarse approximations of conformational free energies. Further, we demonstrate that AF2 can predict the functional consequences of mutations. We discuss the relevance of our findings in the context of the evolving landscape of biomolecular modeling and simulation.

### Theory

Collective variables (CVs) are typically defined as a set of variables that describe the collective behavior of groups of atoms or residues. These variables capture essential aspects of protein dynamics, such as global motions, local conformational changes, and interactions between different regions. By focusing on a reduced set of collective variables instead of considering the full atomistic details, the partition function governing key aspects of a protein’s dynamics can be more effectively sampled.

More formally, one can consider a classical biomolecular system with Hamiltonian

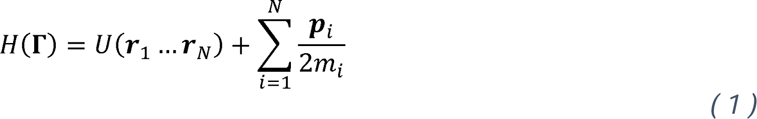

where Γ = (***r***_1_ … ***r****_N_*, ***p***_1_ … ***p***_*N*_) defines the collection of coordinates and momenta, respectively, of all particles in the system. The partition function, *Q*, is given by

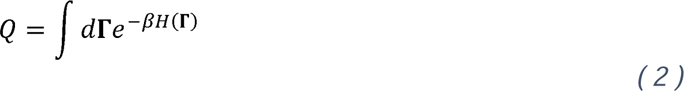

such that the probability of conformation **Γ***_k_* is

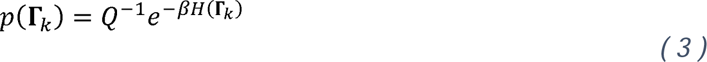

Here, *β* ≡ 1/*k_B_T*, where *k_B_* is the Boltzmann constant and *T*, is the temperature.

In practice, sampling states to approximate Q is computationally intractable for biomolecular systems. Instead, we identify a set of CVs, **ξ** = (ξ_1_, ξ_2_, …, ξ*_m_*), that describe the relevant biological motions and can be calculated from **Γ**. To analyze some observable as a function of a set of CVs, *A*(**ξ**), from the discrete trajectory frames of an MD or Markov chain Monte Carlo (MCMC) simulation, we would compute the ensemble averaged observable as

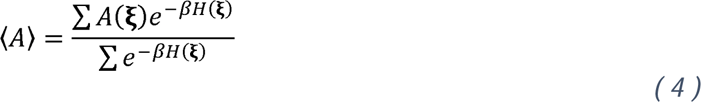

Contact distances are frequently utilized as CVs to describe conformational changes in biomolecular simulations.

Provided an MSA, AF2 builds a predicted a distance distribution for each Cβ – Cβ (or Cα for glycine residues) in the target protein ^30^. The central hypothesis of this manuscript is that the collective contact distance distributions predicted by AF2 contain relevant information that can approximate Boltzmann distributions provided the relevant conformational states can be adequately described by these contact distances.

Several recent investigations suggest that AF2 learns about the energy surface, and therefore conformational partition function, of its target protein. It has been proposed that AF2 utilizes the coevolutionary information within a target protein MSA to identify a neighborhood of plausible structures before locally moving toward a single output structure using its learned representation of model accuracy ^31^. Similarly, recent investigations have demonstrated that reducing the depth of the MSA in an AF2 protein structure prediction task can cause AF2 to generate additional native-like conformers spanning the dynamic range of that protein, but that these structures are often predicted with lower confidence ^32,33^. In the present manuscript, we take these observations one step further and explore whether thermodynamic information regarding protein dynamics can be derived from AF2 predictions.

## Results

### Correlation of AlphaFold2 contact distance distributions with distributions obtained from EPR spectroscopy

For a given protein system, AF2 learns a Cβ – Cβ (or Cα for glycine) contact distribution discretized between 2.3125 and 21.6875 Å at regular 0.3125 Å intervals, with one additional bin for the probability of a distance being greater than 21.6875 Å for a total of 64 bins. EPR spectroscopy is one of only a few experimental methods that provides contact distance probability distributions. Therefore, we first sought to compare distance distributions obtained from EPR spectroscopy to the distributions predicted for Cβ – Cβ pairs by AF2. We selected T4 lysozyme as our model system because it has been well-described at these shorter distances using continuous wave EPR spectroscopy ^34,35^.

We identified six contacts between 12 unique residues in T4 lysozyme (Figure 1A). For each of these contacts, we retrieved the distogram logits from AF2 and computed contact distance probabilities (see **Methods**). We compared these probabilities against the EPR distance distributions for the same contacts (Figure 1B – G). Interestingly, we observe similar relative probabilities among the contact distances for AF2 and EPR, such that lower probability distances in AF2 are generally lower probability distances by EPR spectroscopy. Although theAF2 distributions are broader, the overall agreement between the AF2 predictions and experiment suggests AF2 distance distributions may relate to protein conformational dynamics.

**Figure 1.**
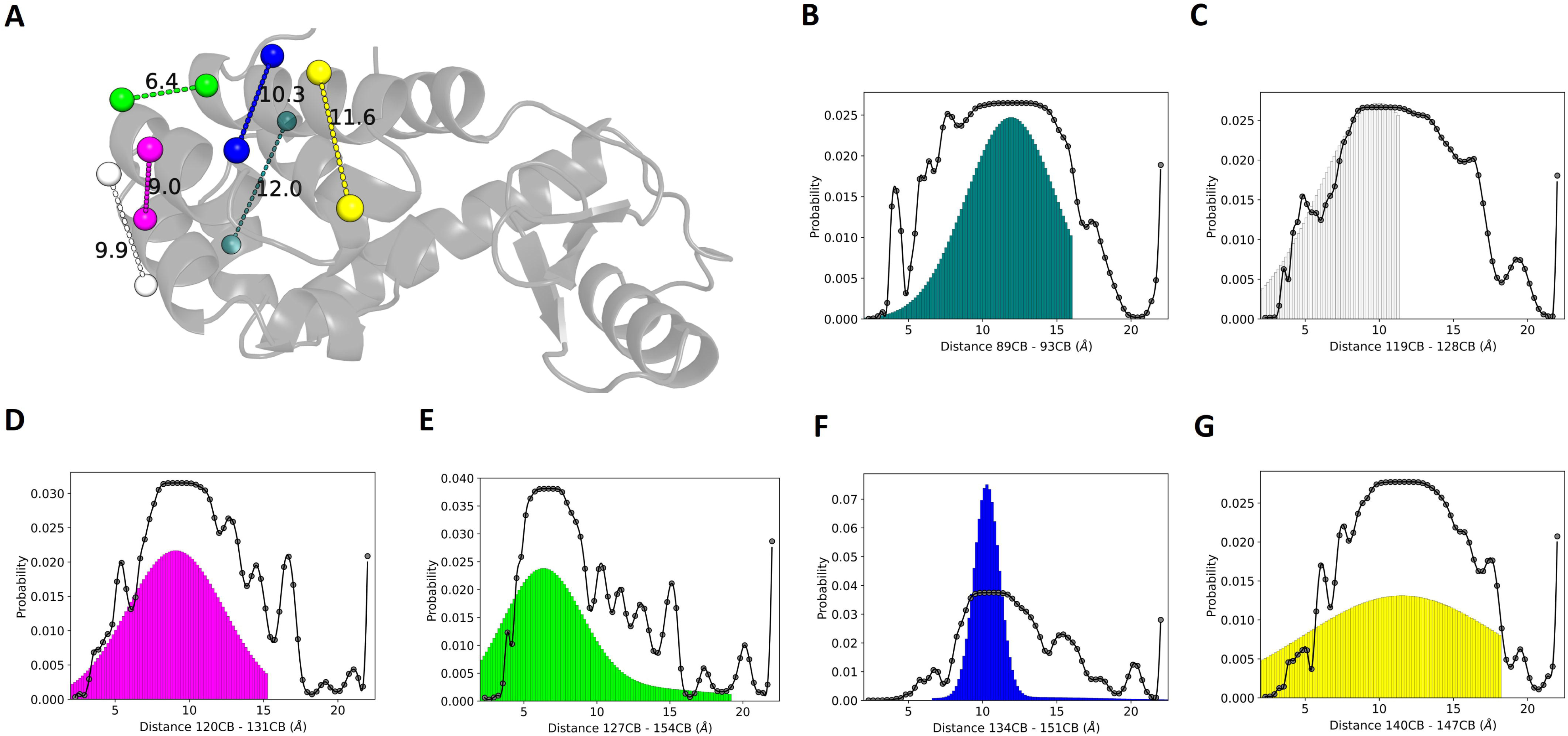
Comparison of AF2 predicted contact distance probabilities with EPR spin-label distance probability distributions in T4 lysozyme. (A) Structure of T4 lysozyme. Tertiary structure illustrated in cartoon format (light blue, semi-transparent). Cβ atoms shown as spheres. Cβ atom distances between residues 89 and 93 (teal), 119 and 128 (white), 120 and 131 (magenta), 127 and 154 (green), 134 and 151 (blue), and 140 and 147 (yellow) are represented with dashed lines and labeled with the crystallographic distances in units of Å. (B – G) Predicted Cβ contact distance probabilities from AF2 (black plot lines) and experimental EPR distances probabilities (colored plot histograms). The AF2 probabilities are normalized to sum to 1. The EPR spina label distances are corrected to the crystallographic Cβ distance.

### Reweighting the conformational free energy landscape of EGFR kinase domain obtained from MD simulation with AlphaFold2 contact distance probabilities

In our lysozyme example, the contact distances that we analyzed are found in the well-folded C-terminal domain of the protein. The functional role of T4 lysozyme is not governed by folding/unfolding events or large conformational changes within that domain. In principle, this means that AF2 does not need to be sensitive to the relative probabilities of multiple minima in order to correctly predict the structure. We sought to evaluate the ability of AF2 contact distance probabilities to approximate free energy changes in a protein system with multiple functionally important minima. We chose for our investigation the EGFR kinase domain (KD).

The active state of the KD is characterized by an outwardly displaced activation loop and an inward αC-helix (Figure 2A, B) ^36^. The inactive state is characterized by an inward activation loop that blocks access of peptide substrate from the ATP binding pocket. The αC-helix oscillates between an outward and a partially inward conformation in the inactive state (Figure 2C) ^3,9,36^. Previously, we characterized the conformational free energy landscapes of wild-type EGFR and several oncogenic exon 19 deletion mutations using conventional and biased MD simulations ^3^. Here, we start by re-analyzing our conventional MD simulations of the wild-type EGFR kinase domain.

**Figure 2.**
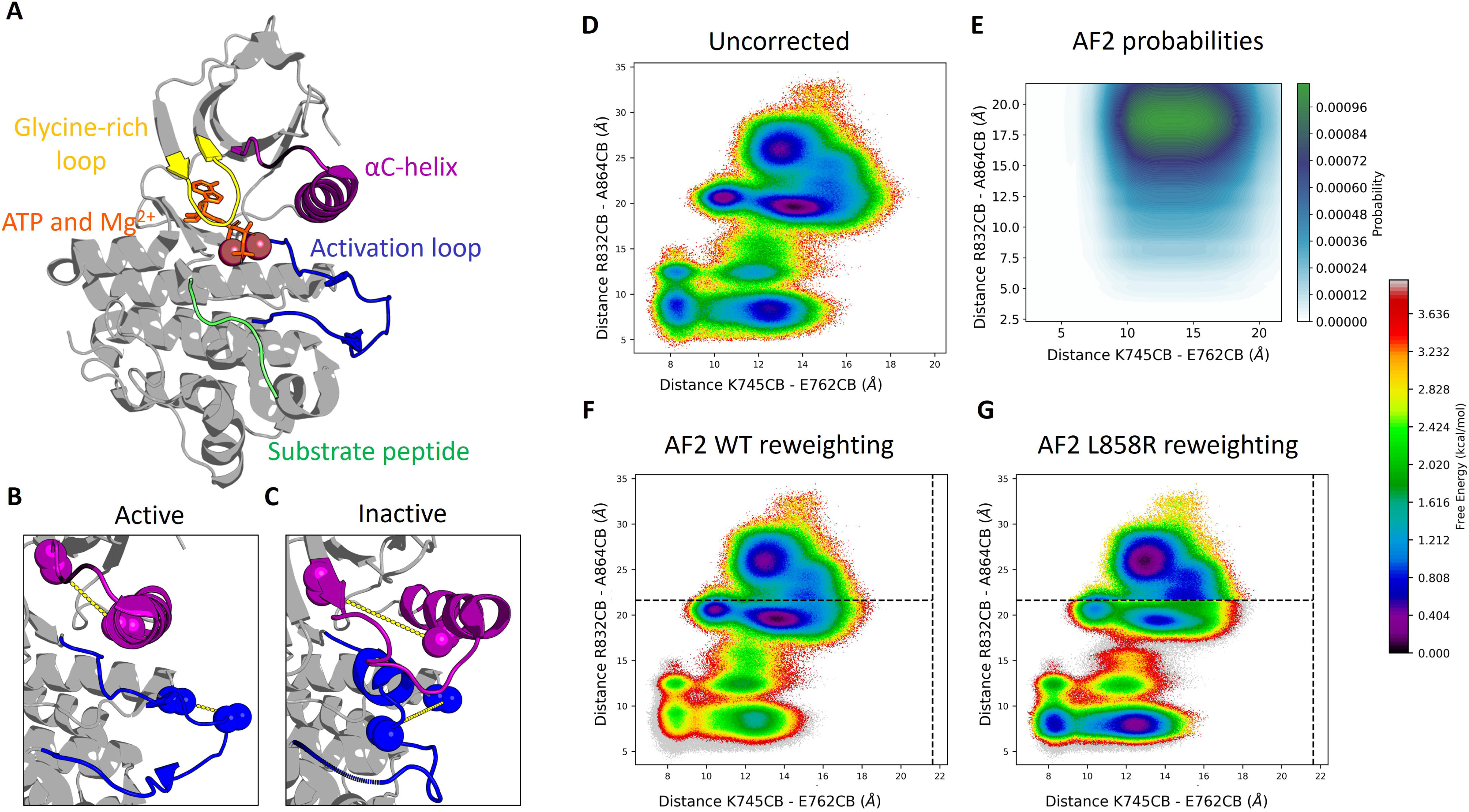
Reweighting MD simulation trajectories with AlphaFold2 probability distributions. (A) Model of EGFR kinase domain in the active state in complex with substrates. Activation loop and αC-helix in the (B) active and (C) inactive states. (D) Conformational free energy landscape of EGFR kinase domain estimated from 6 independent MD simulations started from the active (3×4000 ns) and inactive (3×6000 ns) states. (E) Probabilities obtained from the normalized joint probabilities of the two collective variables defined by the AlphaFold2 contact distance distributions. (F) AF2-reweighted conformational free energy landscape for wild-type and (G) L858R.

We analyzed six independent MD simulations of EGFR KD totaling 30.0 µs (three trials of 6.0 µs each starting from the inactive state and three trials of 4.0 µs each starting from the active state) (Figure S1). Monomeric EGFR KD is known to preferentially adopt the inactive state ^36^; however, the free energy landscape constructed from raw sample state counts of the conventional MD simulations suggest that the active and inactive states are approximately equally likely (< 1.0 kcal/mol difference) (Figure 2D).

We reasoned that if AF2 contact distance probability distributions approximate the actual conformational Boltzmann distribution, then we could use the AF2 probabilities to reweight the incorrect free energy surface, analogous to reweighting with the Markov state model stationary distribution probabilities ^37^. Indeed, our results demonstrate that the EGFR KD activation free energy surface reweighted with AF2 probabilities prefers the inactive state by approximately 1 –2 kcal/mol (Figure 2E, F), which is in agreement with results obtained through umbrella sampling and metadynamics simulations ^3,10,11^. These results further suggest that AF2 contact distance distributions may approximate the likelihoods of unique conformation states of protein systems.

### Sensitivity of AlphaFold2 contact distance probabilities to differential stabilization of folded minima

Mutations in the EGFR kinase domain are a common cause of non-small-cell lung cancer (NSCLC). Oncogenic EGFR mutations alter the native function of the wild-type receptor by changing the relative probabilities of the conformational states along the active-inactive state free energy surface ^9,38^. Wild-type EGFR kinase domain samples space more broadly than its oncogenic mutants ^3,9^. Under the assumption that oncogenic mutants stabilize a subspace of the same conformational free energy surface as described by the CVs, one should in principle be able to obtain the correct mutant free energy surface by reweighting the sampled space. We estimated probabilities for the two CVs for the well-described EGFR L858R oncogenic missense mutation and applied these probabilities to reweight the wild-type EGFR conformational free energy surface. Interestingly, we do correctly observe a shift in free energy favoring occupancy of the active state (Figure 2G). Note, however, that AF2 probabilities at distances beyond 21.6875 Å are unreliable due to the final distogram bin’s use as a catch-all bin (Figure 2F, G; dashed lines).

Mutation sensitivity is typically discussed in one of two contexts: (1) the effect of a mutation on the conformational equilibrium of a protein wherein the equilibrium is between two or more well-folded states (i.e., the ΔΔG^0^ of conformational change), or (2) the effect of a mutation on the thermostability of a protein (i.e., the ΔΔG^0^ of folding). To extend our initial free energy landscape reweighting results, we investigated the first context of mutation sensitivity by analyzing the effects of cancer-causing EGFR mutations on the AF2 contact distance probability distributions. Specifically, we focused on the 832Cβ – 864Cβ distance as a proxy for activation loop motion.

The missense mutations G719S, L747P, and L858R are known activating mutations in EGFR KD. By introducing the mutations across the MSA as previously described, AF2 correctly identifies the shorter contact distance corresponding to the active conformation to be higher probability than the longer contact distance corresponding to the inactive conformation (Figure 3A). As controls, we also generated AF2 contact distance probabilities for EGFR wild-type and a mutant that is known to cause resistance to covalent inhibitors of EGFR without activating the protein, C797S. For both of these latter cases, we see a preference for the inactive conformation (Figure 3A). We also evaluated the predicted active state probabilities for a heterogenous group of activating mutations located in the β3 – αC linker region of EGFR KD collectively referred to as exon 19 deletion mutations ^3^. In total, we computed AF2 contact distance probabilities for 59 unique exon 19 deletion mutations, and all of them were correctly predicted to stabilize the active conformation (Figure 3B).

**Figure 3.**
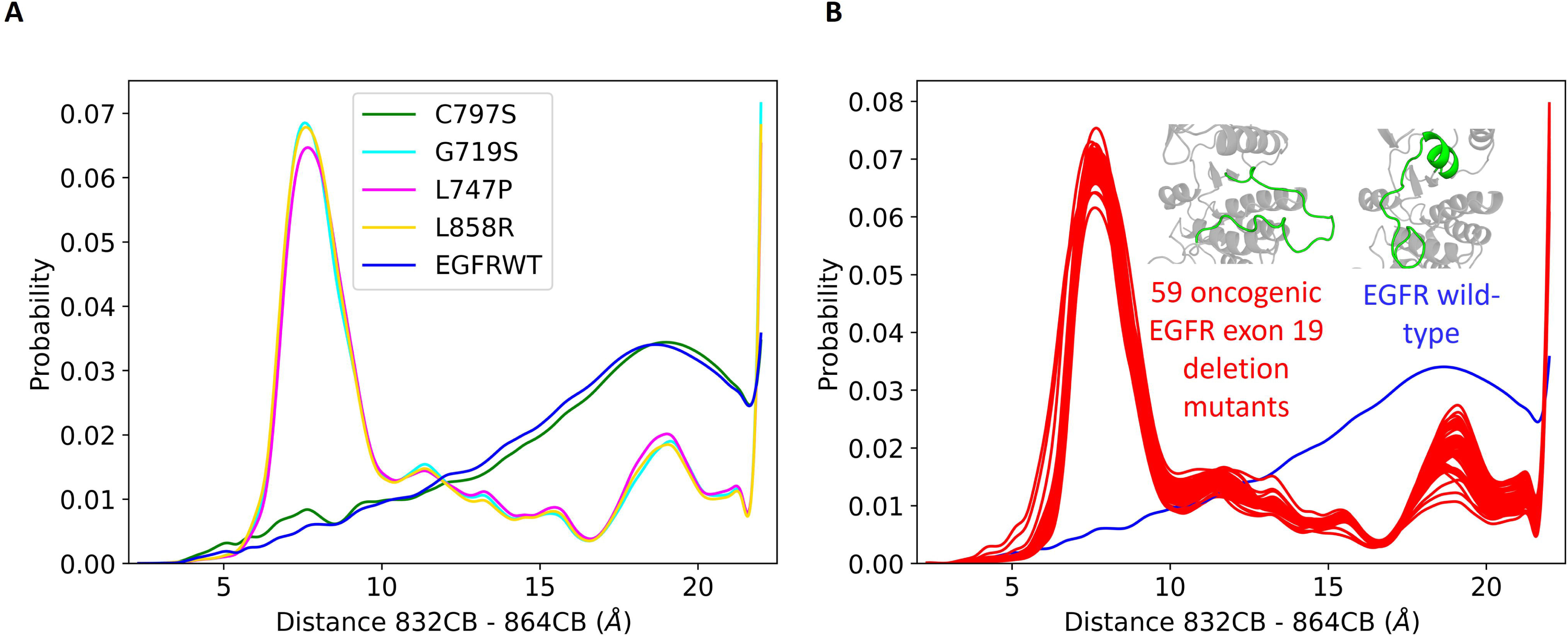
Screening oncogenic mutations by sequence with AlphaFold2 probability distributions to determine oncogenicity. (A) Sequences from EGFR variants: wild-type (blue), the non-activating covalent inhibitor resistance mutation C797S (green), and the activating mutations G719S (cyan), L747P (magenta), and L858R (yellow). Sequence predictions for 60 EGFR kinase domains, wild-type (blue) and 59 known oncogenic exon 19 deletion mutation variants (red). Each sequence was independently provided as input for structure prediction with AlphaFold2. Multiple sequence alignments were independently performed for each variant.

To demonstrate the generality of AF2 mutation sensitivity, we tested the mutation sensitivity of AF2 contact distance probabilities for a series of mutations in the discoidin domain receptor 1 (DDR1). DDR1 binds and is activated by collagen in the extracellular matrix ^39,40^. Unlike the EGFR KD, the DDR1 KD displays slow activation kinetics and preferentially adopts the DFG-out conformation ^41–43^. Hanson and colleagues combined an aggregate *multi-millisecond* set of MD simulations of DDR1 kinase utilizing the distributed computing powers of Folding@home ^44^ with mutagenesis and biochemical analyses to characterize the conformational free energy landscape of the DFG flip (i.e., transition between the “out” and “in” states). They identified and validated the alanine substitutions Y755A and Y759A as mutants that reduce the preference of DDR1 for the DFG-out conformation ^41^. We generated AF2 contact distance probability distributions for DDR1 KD wild-type, Y755A, Y759A, and double Y755A-Y759A. For CV, we used the Cβ distance between the DFG F748 and the back hydrophobic pocket F725, such that longer distances reflect the DFG-out conformation and shorter distances reflect the DFG-in conformation. DDR1 wild-type is predicted to adopt the DFG-out conformation and has a peak CV probability between 16.0 and 18.0 Å (Figure 4A, B). The structure of DDR1 Y755A is also predicted to be in the DFG-out conformation; however, the contact distance probabilities reveal a second peak between 7.5 and 10.0 Å that is approximately equal to the peak between 16.0 and 18.0 Å (Figure 4A, C). Both DDR1 Y759A and Y755A-Y559A are predicted to be in the DFG-in conformation and have progressively increasing peak probabilities in the short distance range and progressive leftward shifts in both short-and long-distance peaks (Figures 4A, D, E). Together with our EGFR results (Figure 3), these data suggest that when suitable CVs are available for a protein target of interest, AF2 contact distance probabilities may be useful as a screening tool prior to more expensive simulations.

**Figure 4.**
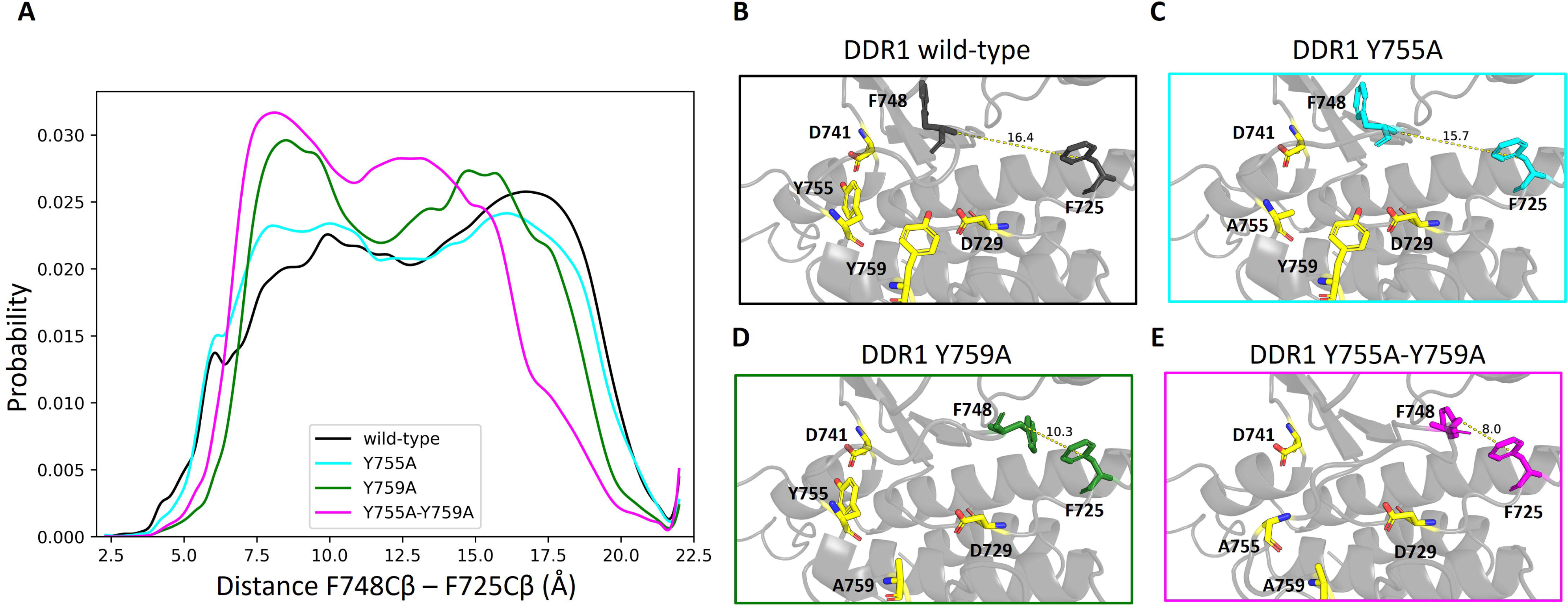
Predicting DFG-motif conformational stability DDR1 kinase using AlphaFold2-derived Boltzmann distributions. Sequences from DDR1 kinase domains: wild-type (black), Y755A (cyan), Y759A (green), and the double mutant Y755A-Y759A (magenta). Each sequence was independently provided as input for structure prediction with AlphaFold2. Multiple sequence alignments were independently performed for each variant. The contact distance probability representing the DFG-in vs. DFG-out conformational transition of the DFG-motif was extracted from each predicted distogram and normalized such that the contact distance probabilities for each mutant sum to 1. The discrete contact distances were interpolated with a cubic spline to yield the curves depicted here.

### Sensitivity of AlphaFold2 contact distance probabilities to thermodynamically destabilizing mutations

Using our new approach, we examined recent findings suggesting that AF2 structure prediction may not be sensitive to sequence variants impacting thermostability ^45,46^. For example, Buel & Walters demonstrate that AF2 predictions of UBA1 ^47,48^ wild-type and L198A yield the same structure despite the latter being a functionally deleterious mutation ^46^. In our hands, AF2 also predicts UBA1 wild-type and L198A to be approximately identical in structure with largely identical predicted probabilities for hydrophobic core residue contact distances to the mutated residue. We also performed three independent MD simulations for UBA1 wild-type or L198A of 1.0 µs each. Even in that short timeframe, the MD simulations suggest that L198A has weakened contacts within the hydrophobic core of the three-helical bundle of UBA1 (Figure 5A, B).

**Figure 5.**
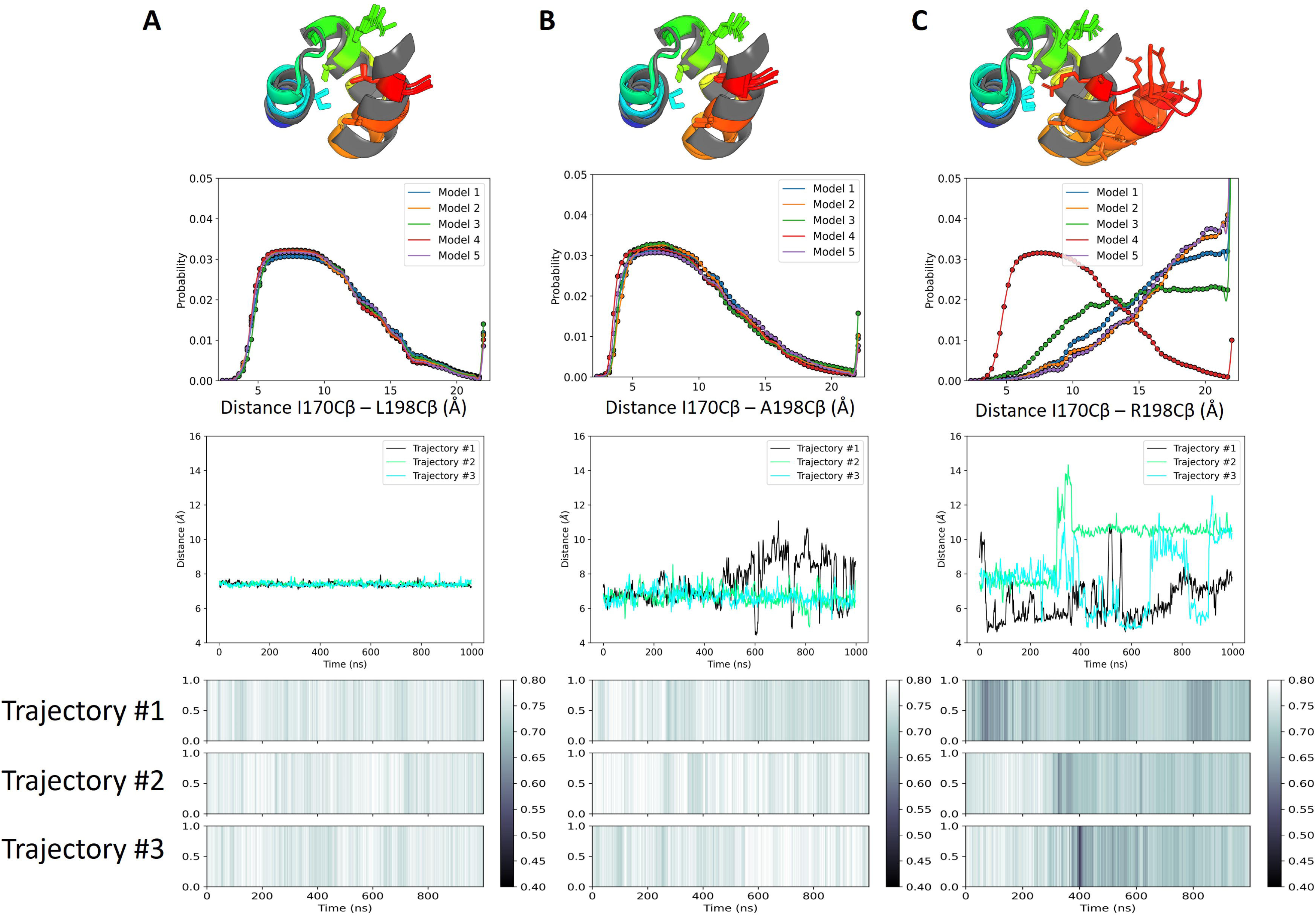
Predicting thermostability of UBA1 mutants. UBA1 (A) wild-type, (B) L198A, and (C) L198R structure (first row), AF2 distance probability distribution for representative interface contact (second row), time-dependent distance distribution of the same representative contact in three independent 1.0 µs all-atom explicit solvent MD simulations (third row), and time-dependent cumulative fraction of α-helix content from the same MD simulations.

We reasoned that a different mutation, L198R, would be more structurally disruptive than L198A owing to its bulky sidechain length and positive charge within the hydrophobic interface. Four out of five AF2 models predicts L198R to have a substantially displaced helix and corresponding higher probabilities of larger distances between the Cβ atoms of interface residues and R198 compared to structures of wild-type and L198A. MD simulations of UBA1 L198R initiated from the structure of wild-type UBA1 quickly become destabilized, and show more rapid loss of α- helical content than wild-type or L198A and Cβ distances more consistent with the AF2 prediction of UBA1 L198R probability distributions (Figure 5C). Our results suggest that AF2 displays mutation sensitivity in its predictions of folding thermostability, but that it is likely less sensitive to physics-based and experimental methods. Consistent with our results is emerging evidence on a larger cohort of proteins suggests that AF2 may have some capacity for predicting folding thermostability ^49^.

### Sensitivity of AlphaFold2 contact distance probabilities to protein-protein interactions

Finally, we tested whether AF2 predicted contact distance probability distributions are sensitive to changes in conformational thermodynamics that occur during protein-protein interactions. As a case study, we analyzed the μ-opioid receptor (μOR). The μOR is a Class A G-protein-coupled receptor (GPCR) whose signaling activation by endogenous or synthetic opioids causes analgesia. ORs are of central interest to the study of pain management and substance use disorder treatment. Class A GPCRs, including μOR, undergo distinct conformational changes to accommodate activation and interaction with a Gα-protein subunit. These changes include a well-characterized large displacement of transmembrane helix 6 (TM6) away from TM3, as well as perturbations to the ligand binding domain ^50,51^.

We predicted the structure of the human μOR 7-transmembrane region alone and in the presence of the human Gαz protein (Figure 6A). The isolated μOR receptor structure adopts an active state conformation despite not being predicted in complex with the Gαz subunit. Overall, the structures of the μOR are virtually identical, differing only modestly (RMSD ∼1.2 Å) with a minor TM6 shift (Figure 6A). Nevertheless, we observe notable differences between predicted contact distance distributions for CVs ^52^ describing the activation mechanism. Specifically, the μOR structure predicted in the presence of Gαz has a steep probability peak at longer TM3 – TM6 distances (∼17.0 – 22.0 Å), which is consistent with the displacement required to bind Gαz. In contrast, μOR structure predicted in the absence of Gαz has a broad TM3 – TM6 distribution (∼10.0 – 22.0 Å) reflective of the shallower equilibrium in the monomeric state (Figure 6B, C). We observe similar changes for TM6 – TM7 extracellular ligand binding domain distance distributions. Importantly, TM6 – TM7 contact distances within the membrane-embedded regions of the helix that are invariant between active and inactive conformations are predicted with the same distribution (Figure 6B, C). We conclude that AF2 contact distance distributions are sensitive to protein – protein interactions.

**Figure 6.**
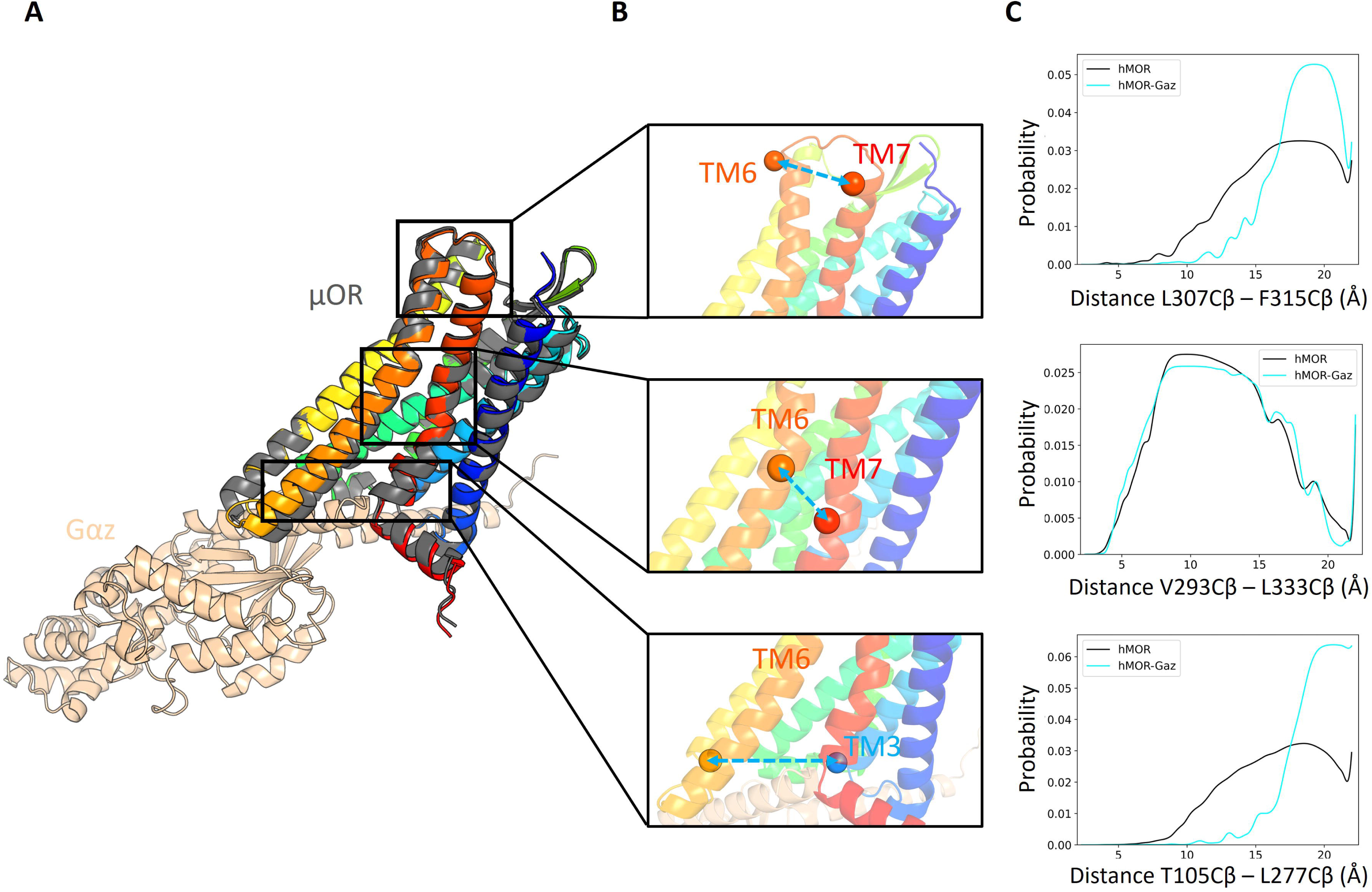
Structure of the human μ-opioid receptor (μOR) predicted in the absence or presence of human Gαz. (A) Structure of the human μOR predicted by AF2 alone (dark grey) or in complex with Gαz (Gαz colored wheat; μOR colored rainbow such that N-terminal residues are blue and C-terminal residues are red). (B) Cβ contact distances that contribute to description of activatiomn mechanism of the receptor. (C) Contact distance probability distributions predicted by AF2.

## Discussion

### Inverting the sampling paradigm with learned partition functions

In traditional physics-based sampling algorithms, such as MD or MCMC simulations, the system’s force field or energy function is used to evaluate the system’s conformation states. By iteratively identifying new conformation states and evaluating them, a partition function is built. This partition function defines the likelihood of any collection of states in the system, allowing for the comparison of relative free energies (Figure 7A). While effective, this approach can be prohibitively computationally expensive.

**Figure 7.**
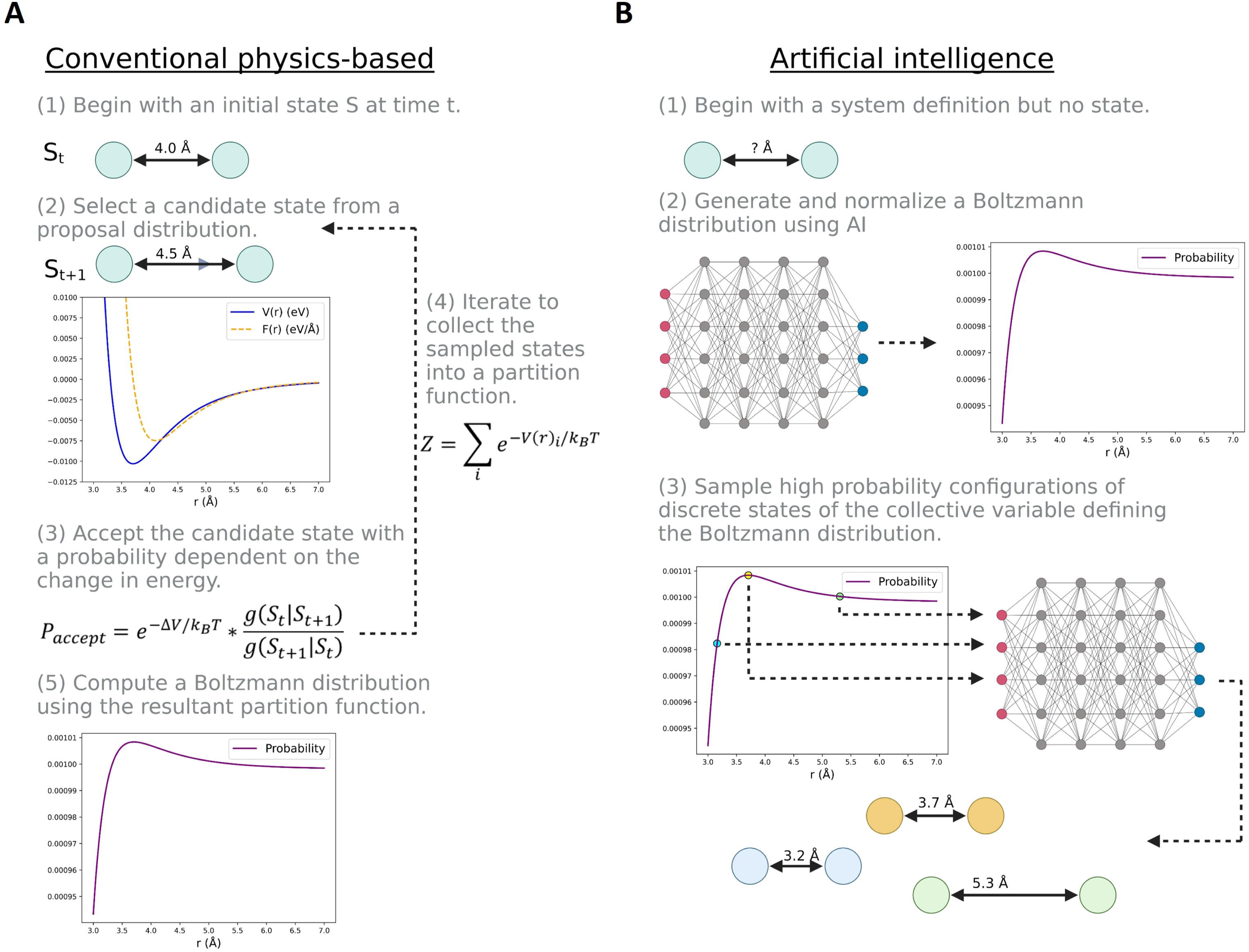
Deep learning methods invert the conventional sampling paradigm by learning a system-specific partition function. (A) In conventional methods, a simulation is progressed in accordance with a physics-based energy function. The collection of states sampled over the course of the simulation define the partition function, which when sufficiently converged enables estimation of the Boltzmann distribution. (B) In some artificial intelligence approaches, distributions of atomic distances, angles, and/or dihedrals are learned for a system of interest. When used as collective variables to describe the functional motions of a system, these distributions effectively enable direct sampling of the relevant free energy surface.

The emergence of AI methods for biomolecular structure prediction are beginning to signal a paradigm shift in sampling of conformational distributions. Instead of sampling discrete states to build the partition function, AI techniques may eventually allow sampling directly from the partition function to identify the most likely conformations. Indeed, in this manuscript we have demonstrated that AF2 can be used in at least some systems to generate coarse approximations of conformational Boltzmann distributions along CVs. With the development of tools such as AlphaLink ^53^, which allows users to pass their own theoretical or experimental contact distance distributions or restraints to OpenFold (an open-source community reproduction of DeepMind’s AF2 ^54^), one can in principal sample the most likely overall protein structures along discrete points of a free energy surface defined by relevant contact distances (Figure 7B). We anticipate that further advances in this direction will prove transformative for biomolecular modeling of protein dynamics.

### Limitations and future directions

While we are not suggesting that AF2 contact distance probabilities are comparably accurate to physics-based methods, we surmise that the learned contact distance distributions approximate physically relevant energy surfaces and may enable some thermodynamic insight into the biomolecular system. Moreover, our results demonstrate the plausibility of AI learning underlying conformational free energy surfaces.

An important limitation of our AF2 conformational free energy landscape reweighting (Figure 2) and oncogenic mutant prediction (Figure 3) results are the boundaries of the AF2 contact distance prediction bins. AF2 was only trained to learn contact distance probabilities between 2.3125 and 21.6875 Å, which for large motions often results in the terminal “catch-all” distance bin having a markedly higher probability than the immediately preceding bins. Thus, a natural extension of our investigations would be to retrain OpenFold to incorporate more distance bins and greater resolution for large dynamical changes.

Finally, our work has implications for the design of proteins with to more realistically account for dynamics modes. Designed proteins, presumably by virtue of their ideal geometries and optimized contacts, are more thermostable and less dynamic than naturally occurring proteins^55^. Our results suggest that designed proteins could be tuned to be more like their naturally occurring counterparts by including metrics for biologically relevant conformational changes using approaches such as those described herein.

## Materials and Methods

### Generation of AlphaFold2 predicted models and contact distance probability distributions

We performed AF2 predictions for each protein system described in the manuscript, which includes T4 lysozyme, wild-type EGFR KD, four missense variants of EGFR KD (G719S, L747P, C797S, and L858R), 59 exon 19 deletion variants of EGFR KD ^3^, wild-type DDR1 KD, three activating variants of DDR1 KD (Y755A, Y759A, and Y755A-Y759A), wild-type UBA1, two destabilizing UBA1 variants (L198A and L198R), μOR, and μOR in complex with Gαz. All mutations were introduced across the entire MSA for all AF2 predictions (i.e., a unique MSA was performed for each mutant). Except when otherwise specified, all subsequent analyses were performed on the top ranked model prediction for each system. We extracted AlphaFold2 contact distance logits from the distogram and converted them to probabilities via the relation 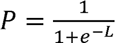 where *P* is probability and *L* is log-odds. Probabilities were then normalized such that 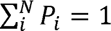 where *N* is the number of discrete contact distances in the AF2 distogram. For 2D AF2 probabilities, normalization occurred over the joint probability space such that ∑*P_i,j_* = 1 where *i* and *j* are distances for two separate contacts.

### Comparison of AlphaFold2 distance distributions to EPR spectroscopy spin label distances

We previously measured methanethiosulfonate nitroxide spin label contact distances with continuous wave EPR spectroscopy for T4 lysozyme ^34^. Spin label distances differ from Cβ – Cβ distances due to the length and flexibility of the sidechain. To enable direct comparison of the Cβ contact distance probability distributions from AF2 with the methanethiosulfonate nitroxide spin label distance probabilities determined by EPR, we adjusted EPR contact distances based on measured Cβ – Cβ distances of the crystallographically determined T4 lysozyme structure. We prepared our reference T4 lysozyme structure by refining a crystallographic structure in Rosetta (PDB ID 2LZM) ^56,57^. We made two mutations to the structure in PDB ID 2LZM, G12R and R137I, to make the sequence consistent with the construct used for EPR distance measurements. Subsequently, we performed a backbone-and sidechain-constrained FastRelax on the structure in Rosetta to yield the final reference structure. For a given spin label distance distribution, each distance bin was offset by the difference between the highest probability spin label distance and the "ground-truth" crystallographic Cβ – Cβ distance.

### Molecular dynamics (MD) simulations

We previously generated 30.0 µs of MD simulations of wild-type EGFR KD across six independent trajectories (three trajectories of 6000 ns starting from the inactive state and three trajectories of 4000 ns starting from the active state) with one of the trajectories transitioning fully from the active to the inactive state ^3^ (Figure S1).

Novel simulation data contributed by the present manuscript include MD simulations for three UBA1 variants: wild-type, L198A, and L198R. The starting structure for UBA1 was obtained from PDB ID 1QZE ^48^. The structures of L198A and L198R were predicted by AF2. Because the structure of L198R diverged from that predicted for wild-type and L198A and we wanted to study the stability of the native fold in the context of the arginine mutation, our starting MD simulation structure for L198R was prepared by simply making the appropriate substitution to wild-type in PyMOL.

Systems were parameterized with Leap in AmberTools22^58^ using the ff19SB force field^59^. Simulations were performed with Amber22 using pmemd.cuda ^58^. Structures were solvated in a rectangular box of OPC explicit solvent neutralized with Joung–Cheatham monovalent ions^60,61^ and buffered on all sides with 15.0 Å solvent. Hydrogen mass repartitioning was performed on solute atoms to allow a simulation timestep of 4.0 fs^62^.

We performed minimization in three stages: (i) The system was minimized with 5000 cycles of steepest descent followed by 5000 steps of conjugate gradient descent (CGD) while protein atoms were restrained with a force constant of 5.0 kcal·mol^−1^·Å^−2^; (ii) the system then underwent 5000 cycles of steepest descent followed by 5000 steps CGD minimization while buffer atoms were restrained with a force constant of 5.0 kcal·mol^−1^·Å^−2^; (iii) all restraints were removed from the system for 1000 steps steepest descent followed by 9000 steps of CGD minimization.

Following minimization, covalent bonds to hydrogen atoms were constrained with the SHAKE^63^ algorithm. Periodic boundary conditions were imposed on the system and the Particle Mesh Ewald (PME) approximation was employed for long-range interactions beyond 9.0 Å. Temperature was controlled using Langevin dynamics with a collision frequency of 5 ps^−1^ during heating and 2 ps^−1^ in production simulations. A unique random seed was used for each Langevin dynamics simulation. Systems were heated from 10K to 100 K in the canonical (NVT; constant number of particles, temperature, and volume) ensemble over 500 ps with a 0.1 fs timestep. Subsequently, systems were heated in the NPT (isothermal-isobaric) ensemble at 1.0 bar with isotropic position scaling from 100 to 310 K over 1000 ps and a 1.0 fs timestep. Pressures were maintained with a Monte Carlo barostat. Production simulations were run in the NPT ensemble at 1.0 bar and 310 K with an integration timestep of 4.0 fs. A total of three independent trajectories for each of the three protein systems were run for 1.0 µs each for a total simulation time of 9.0 µs. Simulation trajectory frames containing solute atoms were collected every 10 ps.

Standard analyses of MD simulations, including time-dependent contact distance and secondary structure content calculations, were performed with CPPTRAJ ^64^. We chose two CVs with which to analyze our MD simulations based on previously published literature ^3,5,8,9,11^ of EGFR KD and adapted them into Cβ contact distances for congruence with AF2: K745Cβ – E762Cβ (αC-helix motion) and 832Cβ – 864Cβ (activation loop motion).

## Acknowledgements

We are grateful to Mohammed AlQuraishi for his insight and feedback.

Benjamin P. Brown is supported by the National Institutes of Health (NIDA DP1DA058349). Jens Meiler is supported by a Humboldt Professorship of the Alexander von Humboldt Foundation and research in the Meiler Lab at Vanderbilt University is supported by the National Institutes of Health (NIDA R01DA046138, NLM R01LM013434, NIA RF1AG06862). Hassane S. Mchaourab and Richard Stein were supported by the National Institutes of Health (NIGMS R01GM077659 and R01GM128087).

The content herein is solely the responsibility of the authors and does not necessarily represent the official views of the National Institutes of Health.

---

**Figure S1. Conventional MD simulations of EGFR kinase domain.** We performed three independent simulations of 4000 ns each starting from the active state (top row). We performed three independent simulations of 6000 ns each starting from the inactive state (bottom row).

