## Supplementary figures and images for "Approximating conformational Boltzmann distributions with AlphaFold2 predictions"

### Supplemental Figure 1

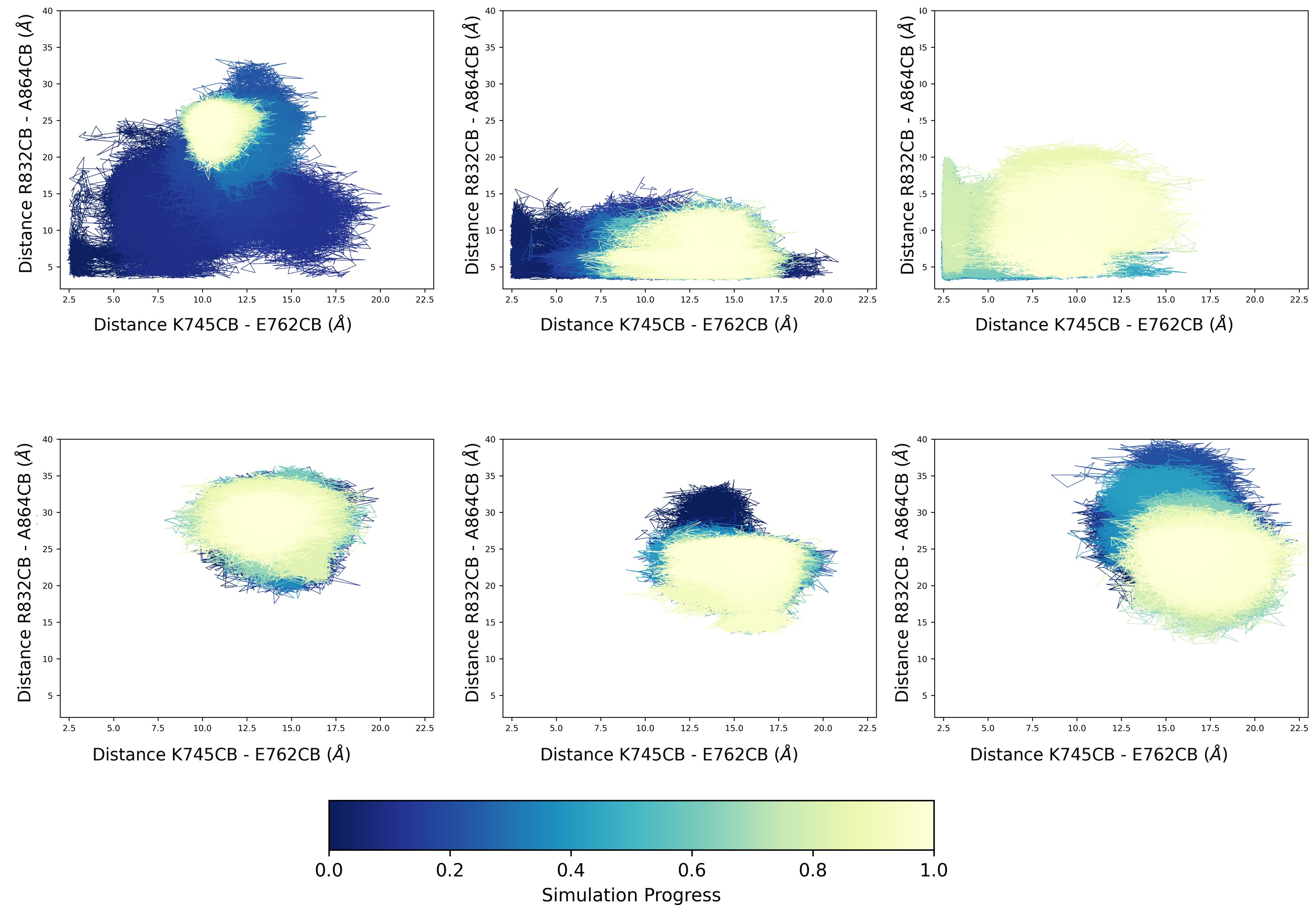
